# Synchrony in Networks of Type 2 Interneurons is More Robust to Noise with Hyperpolarizing Inhibition Compared to Shunting Inhibition in Both the Stochastic Population Oscillator and the Coupled Oscillator Regimes

**DOI:** 10.1101/2023.09.29.560219

**Authors:** Roman Baravalle, Carmen C. Canavier

## Abstract

Synchronization in the gamma band (30-80 Hz) is mediated by PV+ inhibitory interneurons, and evidence is accumulating for the essential role of gamma oscillations in cognition. Oscillations can arise in inhibitory networks via synaptic interactions between individual oscillatory neurons (mean-driven) or via strong recurrent inhibition that destabilizes the stationary background firing rate in the fluctuation-driven balanced state, causing an oscillation in the population firing rate. Previous theoretical work focused on model neurons with Hodgkin’s type 1 excitability (integrators) connected by current-based synapses. Here we show that networks comprised of simple type 2 oscillators (resonators) exhibit a supercritical Hopf bifurcation between synchrony and asynchrony and a gradual transition via cycle skipping from coupled oscillators to stochastic population oscillator, as previously shown for type 1. We extended our analysis to homogeneous networks with conductance rather than current based synapses and found that networks with hyperpolarizing inhibitory synapses were more robust to noise than those with shunting synapses, both in the coupled oscillator and stochastic population oscillator regime. Assuming that reversal potentials are uniformly distributed between shunting and hyperpolarized values, as observed in one experimental study, converting synapses to purely hyperpolarizing favored synchrony in all cases, whereas conversion to purely shunting synapses made synchrony less robust except at very high conductance strengths. In mature neurons the synaptic reversal potential is controlled by chloride cotransporters that control the intracellular concentrations of chloride and bicarbonate ions, suggesting these transporters as a potential therapeutic target to enhance gamma synchrony and cognition.

**Significance Statement:** Brain rhythms in the gamma frequency band (30-80 Hz) depend on the activity of inhibitory interneurons and evidence for a causal role for gamma oscillations in cognitive functions is accumulating. Here we extend previous studies on synchronization mechanisms to interneurons that have an abrupt threshold frequency below which they cannot sustain firing. In addition to current based synapses, we examined inhibitory networks with conductance based synapses. We found that if the reversal potential for inhibition was below the average membrane potential (hyperpolarizing), synchrony was more robust to noise than if the reversal potential was very close to the average potential (shunting). These results have implications for therapies to ameliorate cognitive deficits.

## Introduction

Synchrony in the gamma frequency band is critical to cognition and disrupted in Alzheimer’s disease (Sahu and Tseng 2023) and schizophrenia (Sohal 2022), giving rise to cognitive deficits. Rhythmic stimulation at gamma frequency (Lakatos et al. 2019) is being utilized as a putative therapeutic intervention for cognitive impairment. In order to develop effective and precisely targeted therapeutics, it is imperative to determine the mechanisms underlying gamma synchrony.

Excitable neurons in general can fire action potentials in two modes. One is a pacemaker-like mode in which the input to the neuron is generally above threshold and the firing rate is determined by the mean current input, often called the mean-driven regime in contrast to a fluctuation-driven regime (Petersen and Berg 2016; Schreiber et al. 2009). Networks of coupled oscillatory neurons in the mean-driven regime were discovered to be capable of synchrony mediated by inhibitory synapses (Van Vreeswijk et al. 1994), without the need for excitatory synapses. An influential study showed (Wang and Buzsáki 1996) that inhibitory synchrony was not robust to the levels of heterogeneity thought to characterize physiological networks. In a seminal series of papers (Brunel and Hakim 1999; Brunel and Hansel 2006), Brunel and colleagues analyzed networks of inhibitory neurons and discovered a type of synchrony mediated by inhibition that did not depend on coupling between oscillatory neurons, but instead arose from the interactions between population rate and the synaptic inhibition recruited by that rate. In the mean-driven regime, one would expect the distribution of interspike intervals to be Gaussian. In the fluctuation-driven regime, the neuron is presumed to be in a state with balanced excitation and inhibition (Shadlen and Newsome 1998) biased slightly below the threshold for action potential generation. The balanced excitation and inhibition produce fluctuations that provide a diffusive drive in the presence of a drift back toward the resting membrane potential. This results in a mean-reverting random walk (Uhlenbeck and Ornstein 1930) in the membrane potential (Brunel and Hakim 1999). The firing of individual neurons appears random and the distribution of interspike intervals (ISI) is exponential except for a refractory period. Networks in the fluctuation driven regime produce stochastic population oscillations (SPO) for sufficiently large noise and sufficiently strong inhibition.

The original theoretical work described above used only integrators (Izhikevich 2000) with Hodgkin’s type 1 excitability (Hodgkin 1948), meaning that they can fire at arbitrarily low rates, and studied only current based synapses. The bifurcation structure defines the difference between type 1 and type 2 excitability (Ermentrout 1996; Rinzel and Ermentrout 1998). We have previously shown that PV+ interneurons in layer 2/3 of medial entorhinal cortex exhibit type 2 excitability (Tikidji-Hamburyan et al. 2015), meaning that there is a cutoff frequency below which repetitive firing cannot be sustained. We systematically explored the responses of resonator neurons (Izhikevich 2001) with Hodgkin’s type 2 excitability. We identify two routes to the stochastic population oscillator (SPO), similar to the previously observed routes in type 1 neurons (Brunel and Hansel 2006), except that that type 2 excitability endows neurons with post-inhibitory rebound (Perkel and Mulloney 1974; Rinzel et al. 1998; Tikidji-Hamburyan et al. 2015; Wang and Rinzel 1993). The first route occurs via supercritical Hopf bifurcation from the stationary asynchronous mode, in which the firing rate is approximately constant, into an SPO. The other route is a transition from the coupled oscillator mode to the SPO (Tikidji-Hamburyan et al. 2015). We also studied a biophysically calibrated (Fernandez et al. 2022; Via et al. 2022) model of a network of PV+ interneurons in layer 2/3 of medial entorhinal cortex with conductance rather than current based synapses with either shunting or hyperpolarizing synapses (Vida et al. 2006), or a uniform distribution of reversal potential between the two extremes. Whereas synchrony was more robust in networks with hyperpolarizing synapses compared to shunting or a uniform distribution, the uniform distribution only outperformed shunting below a threshold for synaptic connection strength.

## Methods

### Izhikevich Type 2 model

The equations describing the dynamics of an Izhikevich resonator neuron model are:

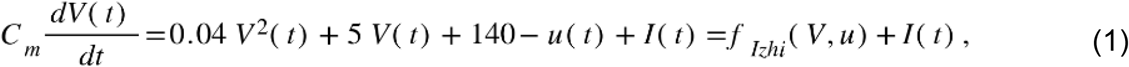

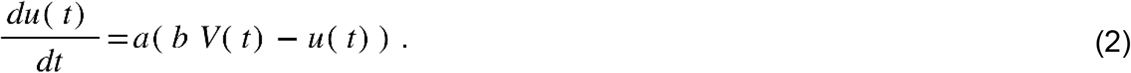

If *V* ≥*v* _peak_, then *V ←c,u* ← *u* +*d* . We use the same values as in our previous work (Tikidji-Hamburyan et al. 2015). The parameters values are a = 0.1 ms^-1^, b = 0.26 nS, c = -65 mV, d = 0 nA. C_m_ is 1 μF/cm^2^. Currents are in nA/cm^2^.

Post-spike adaptation was neglected with parameter *d* set to zero. As the parameter *b* is positive, this model exhibits Type 2 excitability, with a discontinuous *f*-*I* curve.

### Via PV+ fast spiking interneuron model

We also use a calibrated Hodgkin-Huxley type conductance-based model with Type 2 excitability. The model is described in detail in (Via et al. 2022). This single compartment model neurons has five state variables: the membrane potential (V) and four gating variables (m, h, n, and a) that use the same kinetic equations as the original Hodgkin-Huxley model (Baxter et al. 2004; Hodgkin and Huxley 1952), but with different parameters tuned to replicate the dynamics of fast spiking neurons in the medial entorhinal cortex. We included two delayed rectifier K^+^ currents (I_Kv7_ and I_Kv3_). K_V_7 was mislabeled K_V_1 in (Via et al. 2022). The differential equation for the membrane potential (V) of each neuron is

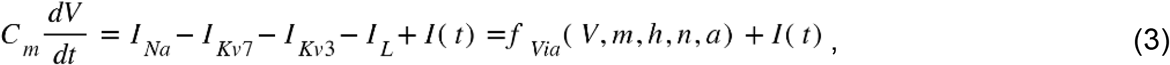

where C_m_=81.4 pF is the membrane capacitance, I_app_ is the external current, I_Na_ is the fast sodium current and I_L_ is the passive leak current. The equations for the intrinsic ionic currents are as follows: 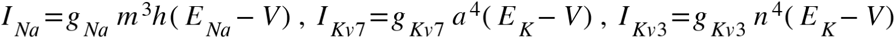,and *I*_*L*_ =*g*_*L*_ (*E*_*L*_ *− V)*, with E_Na_= 50 mV, E_K_= -90 mV, E_L_=-77.8 mV, g_L_=13.8 nS, g_Na_=18929 nS, g_Kv1_=58.5 nS and g_kV7_=784.5 nS. The dynamics of the gating variables are given by *dx* / *dt* = *α*_*x*_ (1− *x*) − *β*_*x*_ *x* for the activation variables (m, n, a) and by *dx* / *dt* = *β*_*x*_ (1− *x*) − *α*_*x*_ *x* for the inactivation variable h, where *α* _*x*_ = *k*_1*x*_ (*θ*_*x*_ −*V*) / (exp((*θ*_*x*_ −*V*) / *σ*_1*x*_) −1) and *β*_*x*_ = *k*_2 *x*_ exp(*V* / *σ* _2*x*_) using parameters in Table 1.

**Table 1.** Parameters for gating variables.

| | $m$ | $h$ | $N$ | $a$ |
| --- | --- | --- | --- | --- |
| $\Theta$ (mV) | -47.95 | -49.72 | 11.32 | 42.85 |
| $\sigma_1$ (mV) | 4 | -20 | 12 | 12 |
| $\sigma_2$ (mV) | -13 | 3.5 | -8.5 | -80 |
| $k_1$ (ms<sup>-1</sup>) | 0.25 | 0.012 | 1 | 1 |
| $k_2$ (ms<sup>-1</sup>) | 0.1 | 0.2 | 0.001 | 0.02 |

### Network definition

We consider a fully connected network of *N* inhibitory neurons. In the subthreshold range (*V* ≤ *V*_*t*_), where *V*_*t*_ is the firing threshold, the membrane potential has the following dynamics:

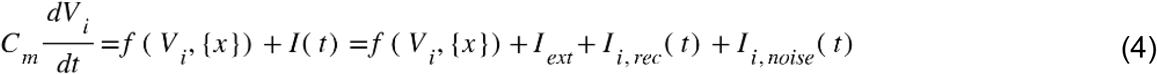

where f(V_i_,{x}) is the function describing the single neuron membrane dynamics, where {x} stands for the gating variables (see Eq. 1 for Izhikevich model and Eq. 3 for Via model), and C_m_ is the membrane capacitance of the model. Here, I_ext_ is a constant external input, I_i,noise_(t) is a noisy external input and I_i,rec_(t) is the recurrent input due to the interactions between the neurons.

The noisy current is modeled as *I*_*i,noise*_ *(t) =σ* η_*i*_(*t*). Here, η_*i*_(*t*) is a white noise uncorrelated from neuron to neuron and from time to time. We define the noise intensity as σ (in nA/cm^2^ for Izhikevich model and nA for the PV+ FS model).

Since the network is fully connected, the recurrent synaptic input is the same for all neurons:

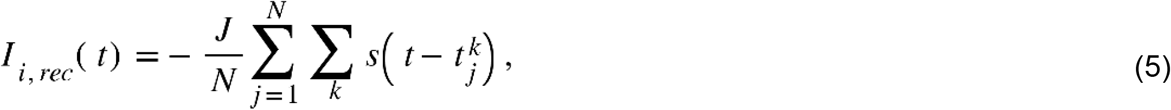

where J is the coupling strength which scales the postsynaptic current for a single synapse (current is given in nA/cm^2^ for Izhikevich model and in nA for Via model), N is the network size and 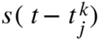 denotes the postsynaptic current (PSC) elicited by a presynaptic spike in neuron j occurring at time 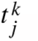.

For the conductance based model, the rhs of Eq. 5 is multiplied by the driving force, current is in nA and J is given in nS

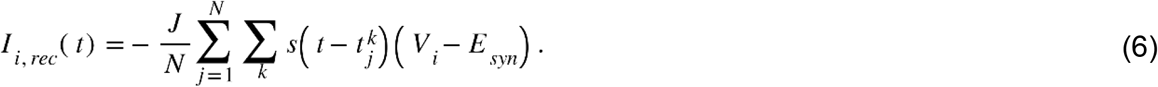

E_syn_ is the synaptic reversal potential. We ran simulations with purely hyperpolarizing (E_syn_=-75 mV) or shunting (E_syn_=-55 mV) synapses, and for the case in which E_syn_ was uniformly distributed between these two values. The first sum is over synapses, whereas the second sum is over spikes. We modeled the PSC function as a biexponential with latency τ_*L*_= 1 ms, rise time constant τ_*R*_ = 1 ms and decay time constant τ_*D*_ = 6 ms.

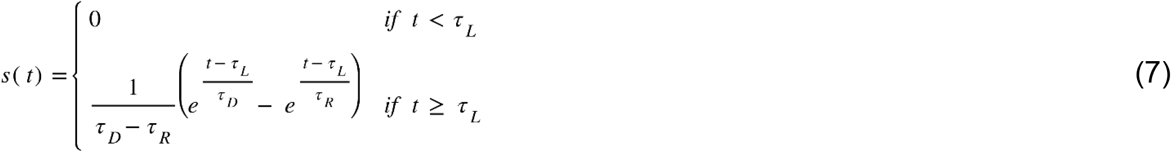

The factor 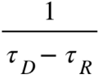 ensures that the integral of the PSC (i.e., the total charge received in the postsynaptic neuron due to a presynaptic spike) is one, i.e., ∫*s*(*t*) *dt=1*. For conductance based synapses the maximum of s(t) is normalized to 1 (Via et al. 2022) *Prediction of Hopf bifurcation surface using mean field theory*.

Brunel and colleagues developed a clever method to find the parameter sets at which an oscillation becomes possible. They used a mean field approximation to the stationary state in which each neuron receives the same oscillation in synaptic current, and each synapse receives action potential inputs at the same frequency as the oscillation in current. In order for an oscillation at a given angular frequency *ω* to exist, the oscillation in the firing rate must have the same phase and amplitude after transformation to an oscillation in current by the biexponential synapse and subsequently back into an oscillation in rate by the individual neurons receiving the oscillatory synaptic input. A linear time invariant system is a cascade of linear operators that scale the amplitude of the input signal via multiplication by|*H* (*ω*)| and shift the phase by ∢ *H* (*ω*) . These quantities can be calculated for a linear system, but the transfer function must be measured at each frequency for a nonlinear system. At a given level of external bias current and noise, the frequency at which the Hopf bifurcation occurs was determined by the value at which all the phase lags around the circuit summed to 0° or 360° or an equivalent multiple. The phase lag for the synaptic latency is ∢ *H*_*L*_ (*ω*) = *ωτ* _*L*_, where *τ* _*L*_ is the latency. The phase lags for the rising and decaying exponentials are ∢ *H*_*R*_ (*ω*) = arctan (−*ωτ* _*R*_) and ∢ *H*_*D*_ (*ω*) = arctan (−*ωτ* _*D*_) where *τ* _*R*_ and *τ* _*D*_ are the rising and decaying time constants respectively. Since the firing rate is positive but the synaptic current is negative, and additional 180° phase lag ensues. The phase lag for the neural model?∢ *H*_*N*_ (*ω*) must be calculated numerically using simulations of a single neuron for each pair of σ and I_ext_ values by applying a sinusoidal input with J=1 then measuring the phase lag of the noisy sinusoidal output. We made predictions at constant firing rate, consistent with previous work (Brunel and Hansel 2006). The constant current I_ext_ was adjusted using a bisection search method, to obtain a constant average firing rate *v*_*0*_ (with an error of 0.5 Hz) for every value of noise intensity σ, coupling strength J and network size N. Since the input is noisy, we found I_ext_ by averaging over 3 seconds of simulation, after discarding the initial 1 second transient.

Next we found the response of the neuron to a sinusoidal input with amplitude 10% of the external bias current to mimic a small sinusoidal perturbation to a steady asynchronous state, with the average frequency fixed at 17 or 30 Hz. Frequency of the input varied between 1 Hz and 250 Hz, with a 2 Hz step. The simulations were run for 50 s and averaged over 20000 realizations. To find the linear gain and the linear phase shift introduced by the neuron between the oscillatory input current to the oscillation in rate output, we used the phase and amplitude of the Discrete Fourier Transform for the peak output frequency.

Finally, the synaptic strength J was adjusted to ensure that the amplitudes of the rate and current waveforms were unchanged from cycle to cycle. The amplitude scale factor for a biexponential synapse with unitary conductance strength is 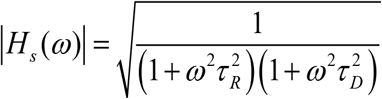. The scaling factor |*H*_*N*_ (*ω*)| for the neural models was calculated using the same method to get the phase lag. Since the connection strength scales the output of the synaptic current, J was set according to 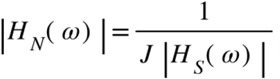.

### Network Simulations

Parameter sweeps were performed on the high-performance computing cluster Tigerfish using BRIAN software (Stimberg et al. 2019). We varied the three parameters J, σ and I_ext_ independently for each of four network sizes (N=800, 1400, 2200, 3000).

### Synchrony measure

We used a synchrony measure to quantify the boundary the boundary between synchrony in our simulations in order to test our mean field predictions. Since synchrony is related to the fluctuations of global variables, it can be defined by averaging these fluctuations over a long time (Ginzburg and Sompolinsky 1994, 1994; Golomb and Rinzel 1993, 1994; Hansel and Sompolinsky 1992). The average membrane potential across a population of neurons at time *t* is 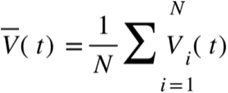. The average over time, or expectation value, of the population-averaged membrane potential is 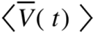. In a completely asynchronous system, any allowable value of V(t) has the same probability for an individual neuron across time as for single neurons across the population. Since these probability distributions are equal, the expectation value of their difference is zero. This makes the expectation value of the population variance 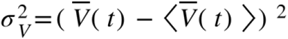 also zero. As a measure of synchrony (Brunel and Hansel 2006), we normalized the population variance to the mean value of the variance 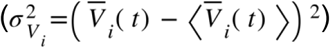 of single-cell membrane potentials V_i_(t):

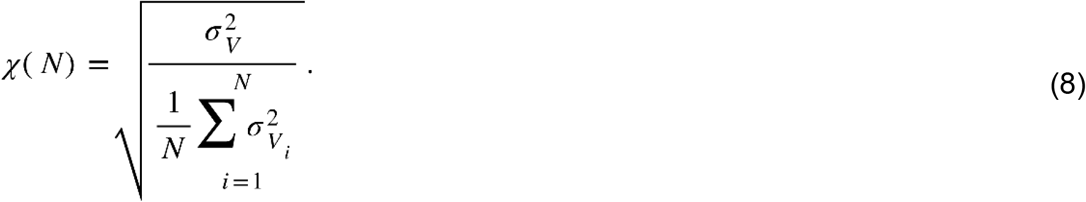

Thus, for a fully asynchronous system χ(*N*)=0. In a completely synchronized system, the expectation value of the population variance is nonzero. Since all neurons have identical activity, the population variance (top term in Eq. 8) is equal to the average expectation value of the individual variance of each neuron (bottom term in Eq. 8), thus χ(*N*)=1. For less than perfect synchrony, the degree of synchrony can be quantified by the measure above. The central limit theorem implies that for an infinite network, *N* →∞, the synchrony measure behaves as

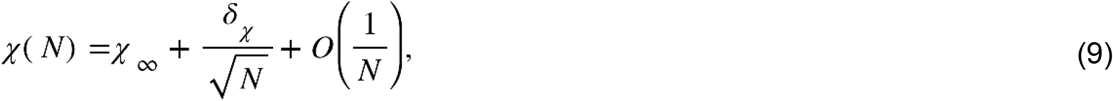

where χ _∞_ is the large *N* limit of χ(*N*) and δ_χ_ measures the finite size correction to χ at the leading order.

For each network size N, we can simulate a network for a given noise intensity, coupling strength and external bias current and calculate the synchrony measure χ(*N*). After simulating several network sizes, we can fit the parameter χ _∞_ using Eq. 9. The set of parameters J, σ and I_ext_ that makes χ _∞_=0 define the Hopf bifurcation manifold, which is the border between synchrony and asynchrony. At each value of J and Iext, we find the critical value of σ_C_ corresponding to the Hopf bifurcation by plotting the values χ_∞_ as a function of σ and noting the point at which χ_∞_ deviates from zero. At constant J and I_ext_, the following mathematical function describes the dependence of *χ*_∞_ on σ:

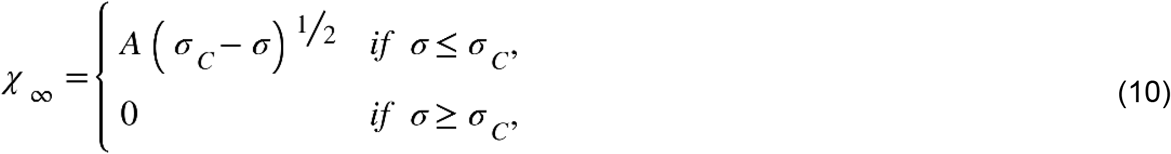

where σ_*C*_is the critical noise at the Hopf bifurcations, for a given J and I_ext_. This method allowed us to find the surface corresponding to the Hopf bifurcation in the 3D space.

### Participation Measure

In order to differentiate between stochastic population oscillators and coupled oscillator regimes we used a method developed by our group based on the cycle by cycle population period, as defined by the peaks of the population histogram (Tikidji-Hamburyan et al. 2015). The spikes were binned in 1 ms windows and a pulse with the height determined by the number of spikes in the bin was placed at the center of the bin. The resultant pulse train was low pass filtered by convolution with a Gaussian kernel with standard deviation of 10 ms and a length of 100 ms, which produced a time series with clear peaks in the network activity to use as a clock to compute the average level of participation in the oscillation, which we track as the average number of spikes per cycle (SPC) normalized by the population size. Note that random peaks of spike rate can be detected in finite networks even in the absence of network oscillations, for example, in random firing or in phase dispersion. In this case, the SPC still reflects the mean firing rate of the population. Values of SPC < 0.6 for a stochastic population oscillator and SPC > 0.9 for coupled oscillator (CO) and post-inhibitory rebound (PIR) regions matched the regions in which an exponential versus a Gaussian distribution of the ISI was observed. Intermediate values corresponded to a transitional oscillatory region with subharmonic peaks in the distribution (for 10 ms bin width). For SPC> 0.9, we distinguished PIR from CO for regions in which the average instantaneous current (external bias plus recurrent input) remained below the Hopf bifurcation at all times.

## Software Availability

https://github.com/RomanB22/NoisyInhibitoryNetwork.git

## Results

### Transitions to SPO in Type 2 model with current based synapses

In order to determine whether our model neurons are in the mean driven or fluctuation driven regime, we first need to understand the bifurcation structure of type 2 neurons that produce hysteresis in the frequency/current relationship. There is a bistable range of injected currents for which either repetitive firing or quiescence can be observed, depending upon whether the external bias current is stepped up such that the current steps become more depolarizing or stepped down. In the Izhikevich type 2 model, there is a subcritical Andronov-Hopf (AH) bifurcation at I=0.2625 nA/cm^2^, and there is a saddle node of periodics (SNP) bifurcation at at I=0.1795 nA/cm^2^. The AH is simply the threshold at which repetitive firing begins as the injected current becomes more depolarizing, and the SNP is simply the threshold below which repetitive firing (minimum value 27 Hz) cannot be sustained as the injected current becomes less depolarizing. The AH and SNP are marked in the next three figures to show that above the AH the neuron is suprathreshold, between the AH and the SNP the neuron is bistable, and below the SNP is it subthreshold. Figure 1 shows the transition from the stationary asynchronous state to the stochastic population oscillator. In the stationary asynchronous rate (Fig. 1A), the mean firing rate of the population is stable over time; an increase (decrease) from the mean firing rate recruits more (less) inhibition, which lowers (raises) the firing rate back towards the mean. At relatively high noise levels, increasing the external bias current triggered a transition through a population Hopf bifurcation to the stochastic population oscillator because the increased bias current increases the firing rate so much that the additional inhibition recruited overshoots the mean rate, initiating an oscillatory cycle of overcorrections (Fig. 1B) with small amplitude near the bifurcation. The oscillatory amplitude grows with distance from the bifurcation (Fig. 1C). The Hopf bifurcation for single neurons described above is unrelated to the Hopf bifurcation that occurs at a population level. However, in these subthreshold regimes, the noise overwhelms the intrinsic dynamics so that the excitability type does not matter. The transition in Fig. 1 for networks of type 2 neurons is qualitatively similar to that found for networks of type 1 neurons. The interspike interval histogram remains approximately exponential as the drive to the network is increased (Fig. 1A1, B1 and C1), despite the emergence of an oscillation in the population rate (Fig. 1B2 and C2) and in the average mean current (green curve in Fig. 1B4 and C4). However, peaks at multiples of the oscillation period will always be observed if there is sufficient data and the temporal resolution is sufficiently fine.

**Fig. 1.**
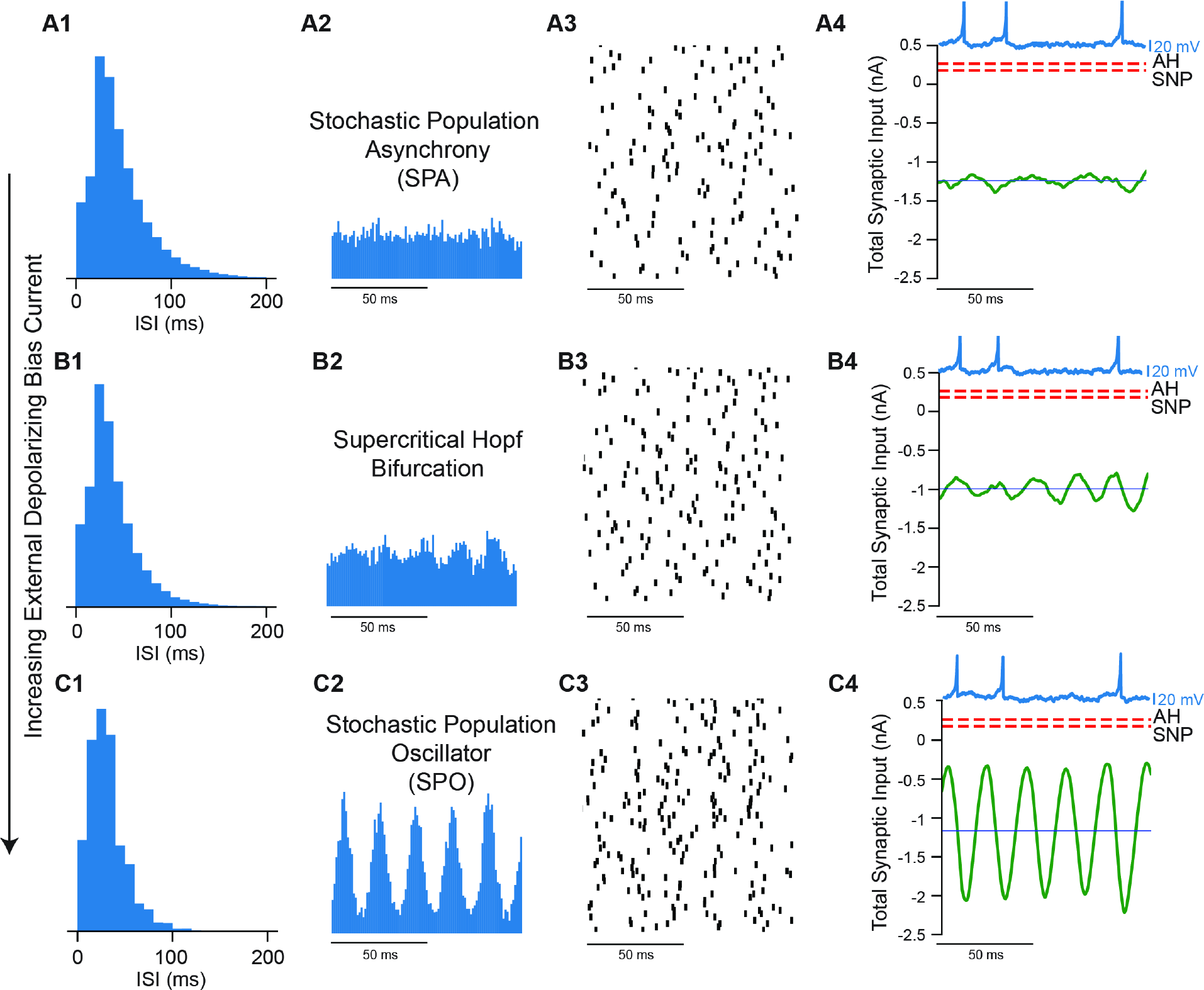
Transition from Stochastic Population Asynchrony to Stochastic Population in a Type II neuron with Increasing Bias Current. **A**. Stochastic Population Asynchrony (I_ext_=0.4 nA). A1. ISI histogram. A2. Spike time histogram. A3. Raster plot (down sampled 71 neurons out of 3000). A4. Current relative to bifurcations (red dashed lines, AH Andropov-Hopf, SNP saddle node of periodics, total synaptic input current averaged over the network (green), and mean current for single neurons (dark blue). Single neuron voltage trace (light blue). **B**. Emergence of stochastic population oscillator at the Hopf bifurcation (I_ext_=1 nA). B1-4 as in A1-4. Spikes Per Cycle (SPC) is 0.49. **C**. Beyond the Hopf, the oscillation amplitude increases (I_ext_= 1.6 nA). C1-4 as in A1-4. In all the three cases the synaptic strength is 7.94 nA and the noise intensity is 3.16 nA. SPC is 0.59. All units are given per cm^2^.

Figure 2 illustrates a different route to the stochastic population oscillator. In contrasts to the transition shown in the previous figure, this transition is gradual and does not involve a bifurcation. Fig. 2A shows a coupled oscillator regime in which the mean current is well below the threshold for repetitive firing. The concept of a spiking threshold does not strictly apply to neurons operating near a subcritical Hopf bifurcation. Between the two thresholds (AH and SNP, see methods) there is a bistable region of single neuron dynamics, in which either depolarization or hyperpolarization can trigger spiking because the stable resting potential is surrounded by an unstable limit cycle. Unlike the case for integrator neurons exhibiting type 1 excitability (Ermentrout 1996; Izhikevich 2007; Rinzel and Ermentrout 1998), an inhibitory input can trigger a spike via post-inhibitory rebound (PIR), as shown in Fig. 2A4. Here not only does the average current remain below the bifurcations that determine the threshold for repetitive spiking, but also the instantaneous values of the total input current to a single neuron. Thus, even though this is clearly a coupled oscillator regime, the mean current is not a reliable determinant of whether the firing is “mean-driven” or fluctuation-driven. Other than the PIR nature of the coupled oscillator spiking, the transition to the stochastic population oscillator for networks of type 2 neurons is similar to that found for networks of type 1 neurons. For the coupled oscillator regime, the ISI distribution is a narrow Gaussian (Fig. 2A1), the population is tightly synchronized (Fig. 2A2 and A3) and neurons spike on every cycle (Fig. 2A4). As the noise increases, neurons begin to skip cycles, leading to the transition regime in Fig. 2B, with peaks in the ISI histogram (Fig. 2B1) at subharmonics of the population frequency corresponding to skipped cycles, and a subthreshold oscillation in the membrane potential of individual neurons during a skipped cycle (light blue trace in Fig. 2B4). Once the firing rate of individual neurons becomes sufficiently sparse, the peaks in the ISI histogram merge into a seemingly exponential distribution (Fig. 2C1) and firing in single neurons appears random (Fig. 2C2 and C4) despite a clear oscillation in the population rate (Fig. 2C3). As stated above, there is no bifurcation, only a gradual transition from a coupled oscillator regime to the stochastic population oscillator.

**Fig. 2.**
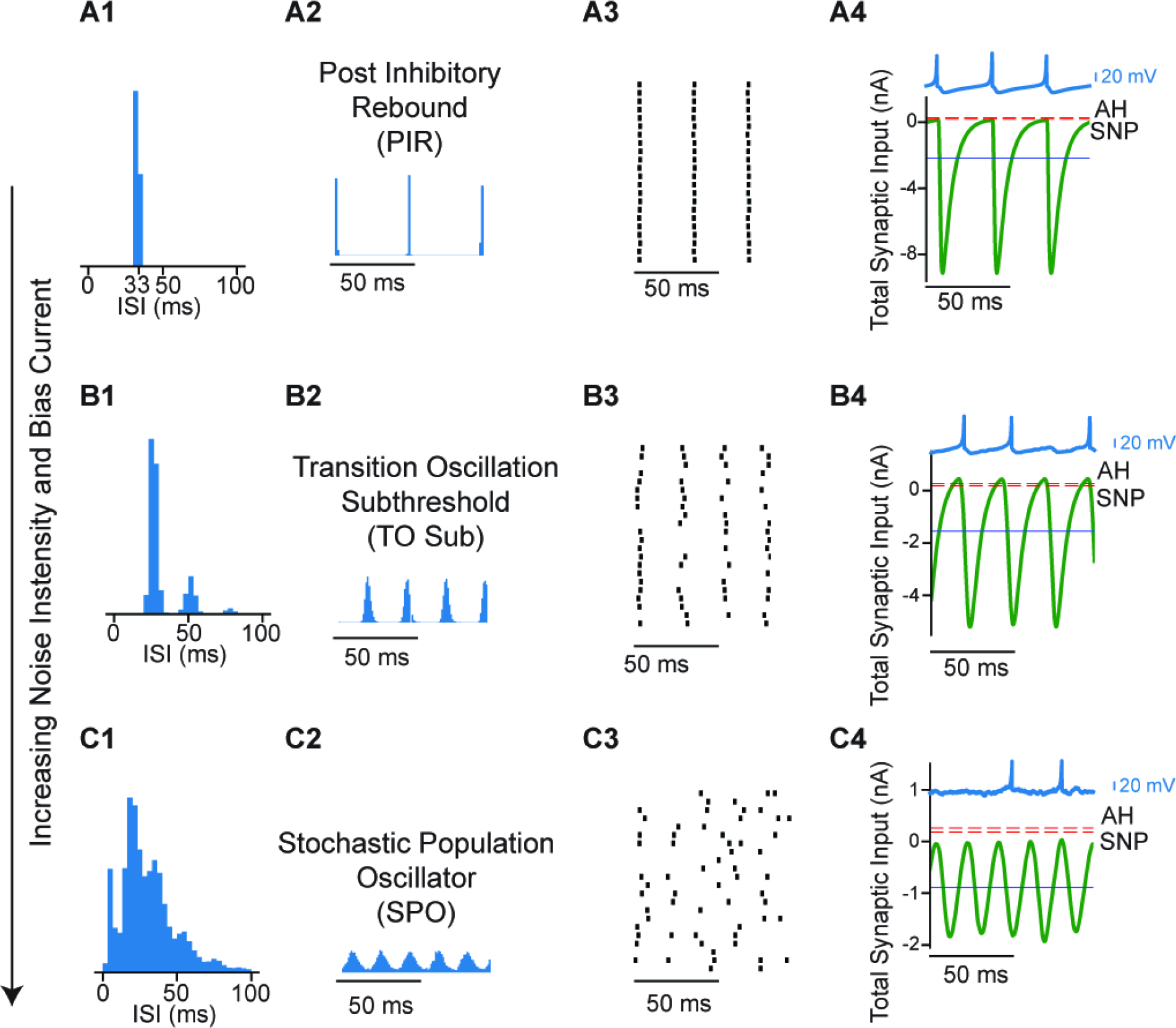
Transition from Coupled Oscillator Synchrony to Stochastic Population Oscillator in a Network of Type II neurons with Increasing Bias Current and Noise Intensity. **A**. Coupled oscillator synchrony due to post-inhibitory rebound (noise intensity is 0.03 nA and bias current is 0.2 nA). A1. ISI histogram. A2. Spike time histogram. A3. Raster plot (down sampled from 3000 to 50). A4. Current relative to bifurcations (red dashed lines, AH Andropov-Hopf, SNP saddle node of periodics, total synaptic input current averaged over the network (green), and mean current for single neurons (dark blue line). Single neuron voltage trace (light blue). Spikes Per Cycle (SPC) is 1. **B**. Gradual transition via cycle skipping (noise intensity is 0.79 nA and bias current is 0.77 nA). B1-4 as in A1-4. SPC is 0.8. **C**. With sufficient noise, the stochastic population oscillator emerges (noise intensity is 3.16 nA and bias current is 1.5 nA). C1-4 as in A1-4. SPC is 0.58. In all the three cases the synaptic strength is 7.94 nA and the mean firing rate is 30 Hz. All units are given per cm^2^.

Fig. 3 shows the transition from asynchrony to synchrony in the coupled oscillator regime. Consistent with previous studies (Brunel et al 2006) and with Figure 1, the transition is via a supercritical Hopf bifurcation. At high levels of noise and external bias current, individual neurons fire irregularly as evidenced by the ISI histogram (Fig. 3A1), the firing rate is stationary over time (Fig 3A2) and the raster plot shows the network is desynchronized (Fig. 3A3). The mean synaptic current (green curve in Fig. 3A4) is above both thresholds for repetitive firing. At the Hopf bifurcation, determined by the mean-field analysis (Fig. 4A), the input to an individual neuron becomes sinusoidal (Fig. 3B4) and remains suprathreshold. In this case, the firing rate was held constant at 30 Hz by decreasing the noise and bias current simultaneously. Decreasing the noise allows the network to start firing in a correlated manner (Fig. 3B3), causing an oscillation in the population rate (Fig. 3B2) that regularizes the ISI histogram, which is now centered around the mean firing rate (Fig. 3B1). As the noise (and bias current) are decreased further, a tight global synchrony emerges (Fig 3C2 and C3) with neurons firing very regularly, evidenced by the narrow, approximately Gaussian ISI histogram (Fig. 3C1). The original theoretical work (Brunel and Hansel 2006) examined a 2D parameter space of the standard deviation of the current noise and the strength of an individual current-based inhibitory synapse, converted to units of voltage by multiplying by the membrane resistance (at rest). The firing rate was kept constant by adjusting the mean level of excitatory external bias current as in Fig. 3.

**Fig. 3.**
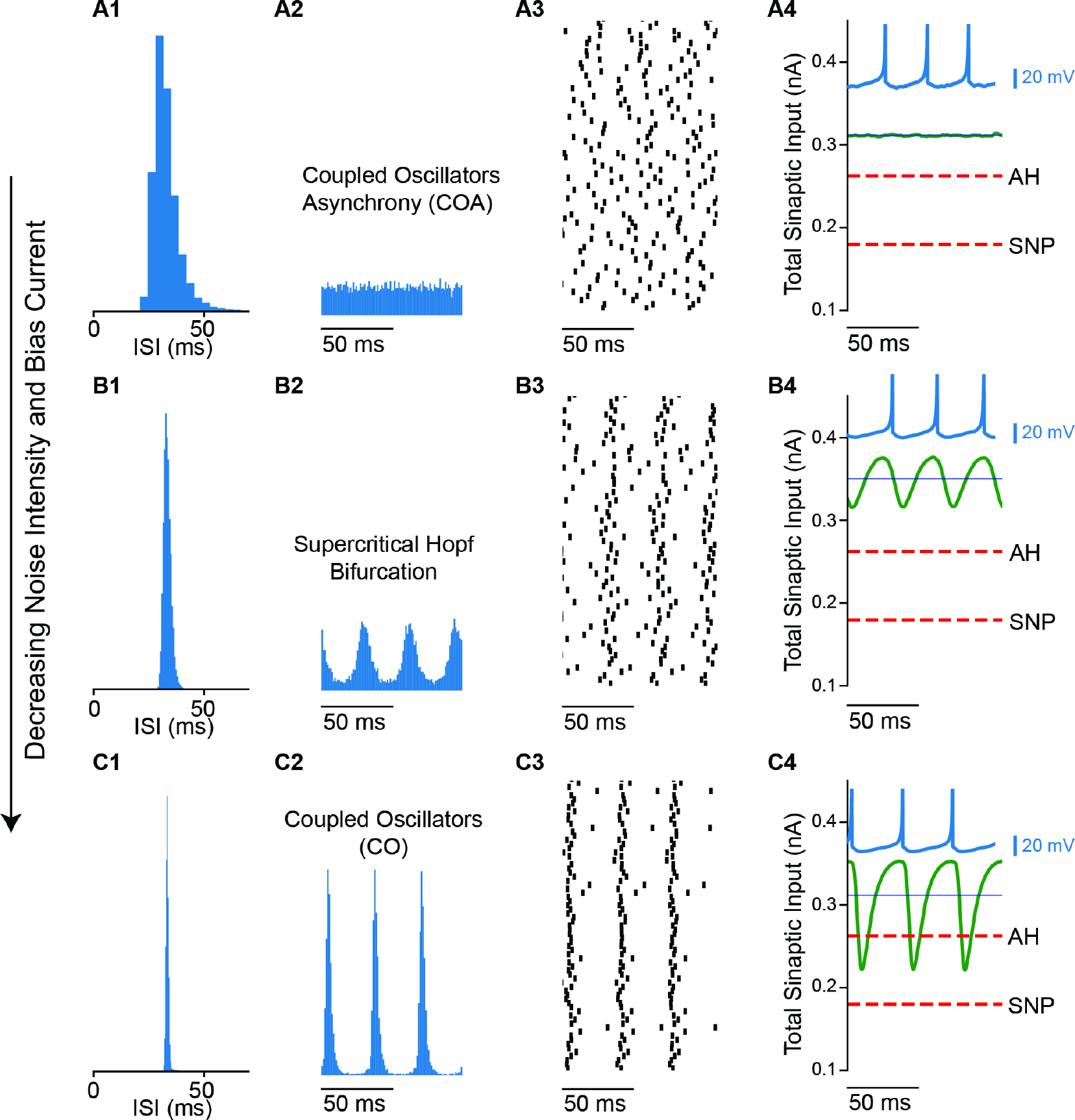
Transition from Coupled Oscillator Asynchrony to Coupled Oscillator Synchrony in a Network of Type II neurons with Decreasing Noise Intensity and Bias Current. **A**. At high noise, the asynchronous state has a stationary mean firing rate (noise intensity is 0.56 nA and bias current is 0.428 nA). A1. ISI histogram. A2. Spike time histogram. A3. Raster plot (down sampled to 71 out of 3000 neurons). A4. Current relative to threshold. Threshold at the saddle node (SN, red dashes) bifurcation, total synaptic input current averaged over the network (green), and mean current for single neurons (blue line). **B**. Transition from network synchrony to asynchrony (noise intensity is 0.14 nA and bias current is 0.398 nA). B1-4 as in A1-4. Spikes Per Cycle (SPC) is 0.98. **C**. Coupled Oscillator Synchrony (noise intensity is 0.05 nA and bias current is 0.359 nA). C1-4 as in A1-4. SPC is 0.99. In all the three cases the synaptic strength is 0.16 nA and mean firing rate is 30 Hz. All units are per cm^2^.

**Fig. 4.**
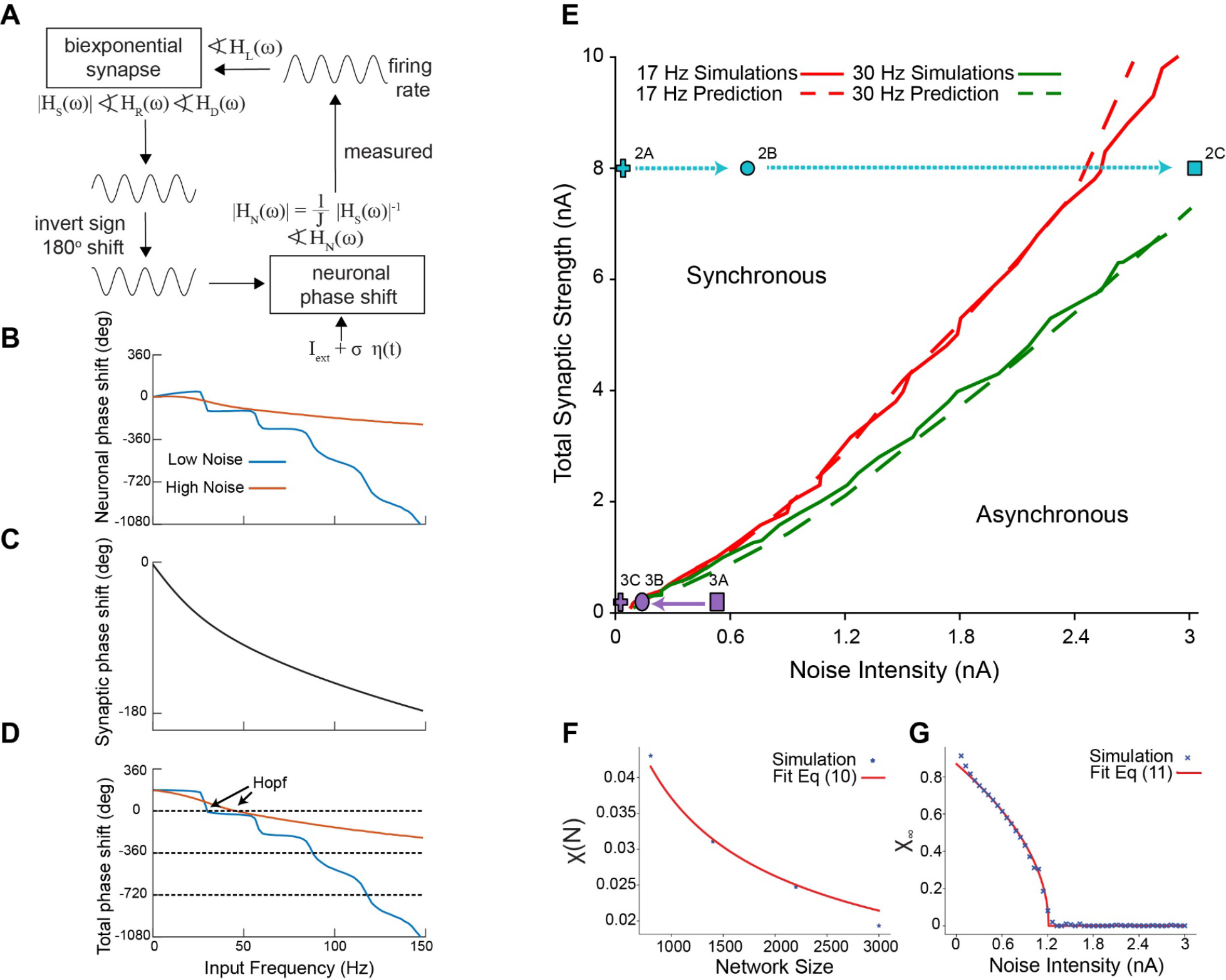
Hopf bifurcation between stochastic asynchrony and synchrony. **A:** Self-consistent criterion for an oscillation in population rate. The sum of the phase shifts must equal a multiple of 360o. **B-D:** Self-consistent phasic relationship for an oscillation in population rate. **B**. Numerically calculated neuronal phase shifts with σ = 0.16 nA, for low noise and σ = 2.8 nA for high noise. Iext is adjusted to maintain a constant firing rate of 30 Hz. **C**. Analytically calculated synaptic phase shift. **D**. The sum of the phase shifts must equal zero or a multiple of 360o (see Fig. 2) **E**. Projection of two curves onto the noise versus synaptic strength plane. Prediction, simulation result. The points corresponding to Figs. 2 and 3 are depicted in cyan and purple, respectively. Dashed arrows depict transition in the synchronous region, full arrows correspond to asynchronous region. **F-G**. Finite-size scaling to find the Hopf bifurcation for Izhikevich model. **F**. Value of χ(*N*) for the four sizes of the network together with the fit of Eq. 9. This fit gives a value of χ_∞_ for each J, σ and I_ext_. In this case J= 2 nA, σ=2 nA and I_ext_=2 nA. **G**. After finding χ_∞_, we find the noise value that desynchronizes the asynchronous state for each value of J and I_ext_. Thus, we end up with a set of values (J, I_ext_, σ_*C*_) which determines the Hopf bifurcation surface. J and I_ext_ are as in F. In E, we

Figure 4 explains how the transitions between asynchrony and synchrony can be predicted using mean field theory, which assumes all neurons in the network receive identical input. The oscillations in firing rate and oscillations in synaptic current give the output and input of the network respectively, and must be self-consistent as illustrated in Fig. 4A and described in the Methods. Figure 4B-D gives an example of the determination of the location of the Hopf bifurcation that gives rise to a stochastic population oscillation (see Methods for detailed explanation). The neuronal phase shift for a low noise case (blue curve in Fig. 4B) and a high noise case (red curve) was calculated numerically by applying a sinusoidal drive to a single neuron (Fig. 4A) with different contributions of a steady bias current and added noise to give the same mean firing rate of 30 Hz. The synaptic phase shift (Fig. 4C) for a biexponential synapse is the same for both cases. The 180o phase shift for the sign inversion shown in Fig. 4A was combined with other two phase shifts in Fig. 4D. An oscillation can arise when it remains in phase with itself after traversing the loop in Fig. 4A, meaning that the sum of the phase shifts totals zero or 360o, or a multiple thereof. For the high noise case there was a single intersection at 46.7 Hz. For the low noise case there were several intersections of the sum of the phase lags with zero phase or multiples of 360o, but only the first intersection at 31 Hz corresponds to a Hopf bifurcation for global synchrony (the other intersections are cluster solutions (Brunel and Hansel 2006)). Figure 4E shows a spot check of the mean field predictions (dashed curves) of the location of the Hopf bifurcation at two constant firing rates, (17 Hz and 30 Hz), projected onto the plane of synaptic strength and standard deviation of the noise. The neuronal phase shift and the transfer function were calculated from the largest Fourier coefficient (Richardson 2008). The mean field theory (dashed curves) accurately predicts the actual Hopf bifurcation (solid curves) in both the coupled oscillator and stochastic population oscillator regimes. The transitions in Figures 2 and 3 were generated at a constant firing rate of 30 Hz, thus they can be visualized in Fig. 4E. The cyan line shows the path from coupled oscillators to stochastic population oscillator from Fig. 2; all values are to the left of the Hopf bifurcation into asynchrony shown at 30 Hz by the green curve. The purple line shows the transition in Fig. 3 from a mode in which neurons are suprathreshold but asynchronous (coupled oscillator asynchrony COA) to a coupled oscillator (CO) synchronous regime when the green curve is crossed from the left. The solid curves in Fig. 4E were obtained by applying the synchrony measure χ(*N*) for networks of size N in Fig. 4F described in the Methods to find the synchronization index for an infinitely large network χ_∞_. The point at which this index deviates from zero is the Hopf bifurcation (Fig. 4F). Hence this metric (as described in the Methods) was used to determine the full 3D structure of the Hopf bifurcation surface (Fig. 5A) that separates synchrony from asynchrony.

**Fig. 5.**
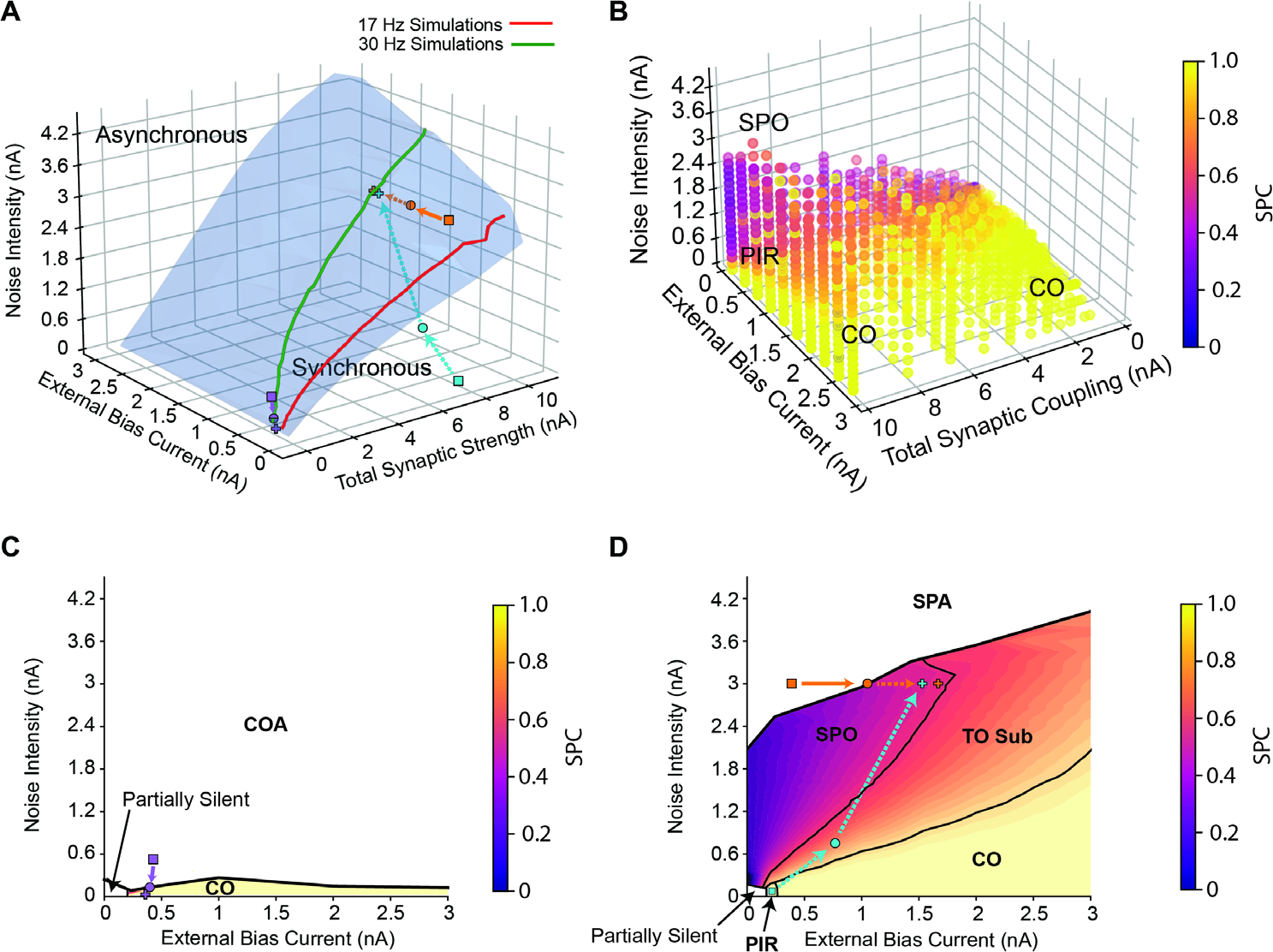
Transitions in the full 3D parameter space. **A**. Bifurcation in the full 3D parameter space. The points corresponding to Figs. 1, 2 and 3 are depicted in orange, cyan and purple, respectively. Dashed arrows depict transition in the synchronous region, full arrows correspond to asynchronous region. **B**. Transition from Coupled Oscillator Synchrony to Stochastic Population Oscillator. The color code indicates the spikes per cycle (SPC), which is the average fraction of oscillations cycles in which an individual neuron spikes. Population oscillations disappear on the upper surface of the colored region, which is the Hopf bifurcation surface from a different perspective than that shown in A. **C**. Plane depicting the low coupling region of the 3D space. The transition is from Coupled Oscillators synchrony (CO) to Coupled Oscillators Asynchrony (COA). Total synaptic coupling is 0.16 nA. **D**. Plane depicting the high coupling region of the 3D space. Total synaptic coupling is 7.94 nA. All units are given per cm^2^.

The transition from the asynchronous stationary state to stochastic population oscillator from Fig. 1 is shown as a purple curve in Fig. 5A and can also be visualized as a downward crossing of the contour of the top of the surface shown in Fig. 5B; that manifold is the Hopf bifurcation shown at a different angle from the one in Fig. 5A.

The transition from coupled oscillators (including the PIR regime) from Fig. 2 to stochastic population oscillator (cyan trace in Fig. 5A) is gradual via cycle skipping. Fig. 5B illustrates the gradual transition via cycle skipping shown in Fig. 2 in the full 3D space as the participation index (SPC) drops from yellow for full participation in which every oscillator participates in each cycle of the population oscillation and decreases gradually (orange) as the stochastic population oscillator region (purple) is approached. The transition from coupled oscillator to stochastic population oscillator (cyan trace) is also shown in the plane section in Fig. 5D; the PIR region is marked for low noise and low bias current, and TO stands for transitional oscillation with subharmonic peaks.

The transition from coupled oscillator asynchrony to synchrony from Fig. 3 is shown in purple in Fig. 5A. The SPC measure is only defined for synchronous regions, thus the coupled oscillator asynchrony and stochastic population asynchrony regions are not evident in Fig. 5B. Figure 5C depicts a plane section of the bifurcation diagram in the low coupling region, and shows the transition in Fig. 3 (purple curve) from a CO asynchronous to a CO synchronous state as the Hopf bifurcation (black curve) is crossed.

### Transitions to SPO in Type 2 model with conductance based synapses are more robust to noise with hyperpolarizing compared to shunting synapses

The results in the previous figures were all obtained with current based synapses. We used conductance-based biexponential synapses and a biologically calibrated model of the PV+ inhibitory interneurons with type 2 excitability in layers 2/3 of the medial entorhinal cortex. Previous studies in the dentate gyrus suggested that the reversal potential of GABA_A_ synapses on PV+ fast spiking basket cells is shunting (Vida et al. 2006) rather than hyperpolarizing. However, a more recent study (Otsu et al. 2020) suggests that in area CA1 the reversal potential can be quite variable. Figure 6A shows the full 3D parameter space and the region of synchrony and asynchrony for both hyperpolarizing and shunting inhibition. Similar to previous results for the coupled oscillator regime (Via et al. 2022), hyperpolarizing synapses are more strongly synchronizing and are thus more robust to noise The red dashed lines in Fig. 6B4 and C4 explain the difference between hyperpolarizing and shunting synapses. Hyperpolarizing synapses cause downward deflections in the membrane potential towards the reversal potential (red dashed line in Fig. 6B4). In contrast, a shunting synapse does not produce appreciable deflections in the membrane potential (reversal potential indicated by red dashed line in Fig. 6C4), but only increases the conductance and resists deflections from that reversal potential. Figure 6B shows a stochastic population oscillation in a network with synapses with a hyperpolarizing reversal potential of -75 mV and exhibiting an approximately exponential ISI histogram (Fig. 6B1), an oscillation in population rate (Fig. 6B2), sparse firing (Fig. 6B3), and a random walk in the membrane potential of individual neurons (Fig. 6B4). We then examined the case in which the reversal potential of the inhibitory synapses was set to a shunting value of -55 mV. A stochastic population oscillator (Fig. 6C) was easily observed in this network. Therefore, the stochastic population oscillator does not require hyperpolarizing inhibition. Nonetheless, the shunting synapses destabilize the asynchronous regime for a lower noise level, which means that the synchrony is less robust to noise for these synapses, both in the coupled oscillator and the stochastic population oscillator regimes.

**Fig. 6.**
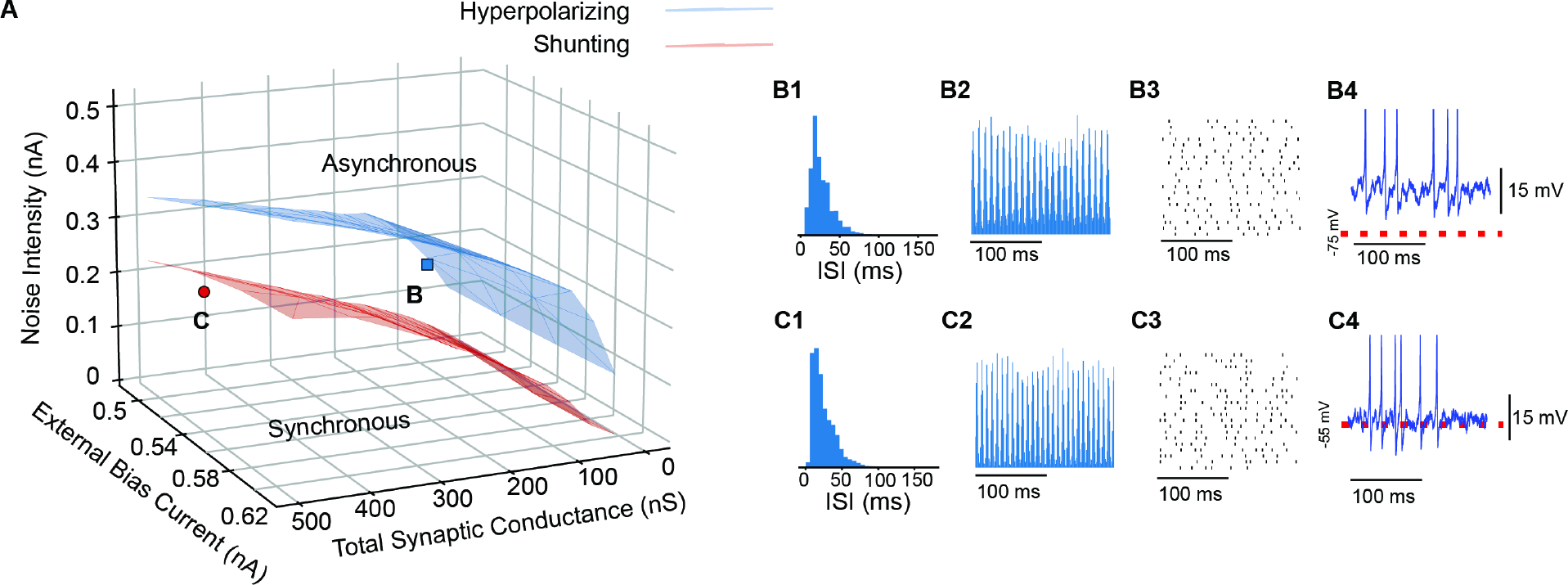
Hyperpolarizing inhibition is more robust to noise than shunting inhibition for all values of coupling and external current. **A**. Bifurcation in the full 3D parameter space for hyperpolarizing (blue) and shunting (red) inhibition **B**. SPO for hyperpolarizing inhibition. Peak conductance is 66 nS. B1. ISI histogram. B2. Spike time histogram. B3. Raster plot (down sampled). B4. Membrane potential of a representative neuron. **C**. SPO for shunting inhibition. Peak conductance is 412.5 nS. B1. ISI histogram. B2. Spike time histogram. B3. Raster plot (down sampled). B4. Membrane potential of a representative neuron. In both cases the noise intensity is 0.16 pA and bias current is 0.5 nA. Red dashed lines indicate the synaptic reversal potential.

### Uniform distribution in reversal potential is less stable than hyperpolarizing for a homogeneous network

According to a recent study, the reversal potential of somatically evoked GABA_A_R-mediated currents for PV+ interneurons in CA1 is distributed from -55 mV to -75 mV, in an approximately uniform way (Otsu et al. 2020). There is also a somatic-dendritic gradient within the PV+ cells, which justifies varying the synapses projecting to a single neuron (Otsu et al. 2020). Thus, we compared networks with either homogeneously hyperpolar-izing or shunting synapses to a network with uniform distribution between the two extremes. Fig. 7 shows that although the hyperpolarizing synapses are still the most synchronizing ones (Fig. 7A and B) across all the parameter space, a uniform combination of shunting and hyperpolarizing synapses is not always more synchronizing than shunting (Fig. 7A and B). For a region with low total synaptic coupling (below around 2 to 2.5 nS) the uniform distribution of reversal potentials is more stabilizing than shunting, whereas the opposite occurs above these values (Fig. 7B).

**Fig. 7.**
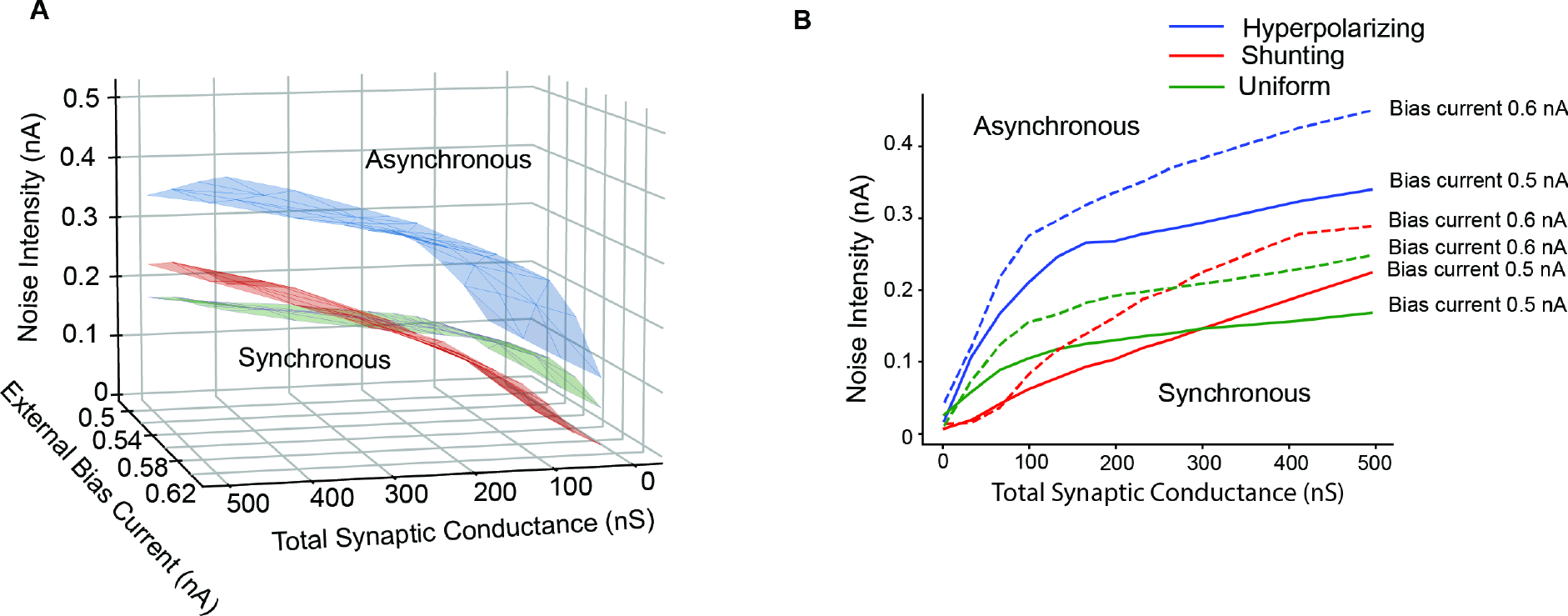
Uniform distribution of reversal potential is less robust than hyperpolarizing for all values of bias current, noise and coupling, and less robust than shunting for high synaptic coupling. **A**. Bifurcation in the full 3D parameter space for hyperpolarizing (blue), shunting (red) inhibition and uniform distribution for reversal potential between -55 and -75 mV. **B**. Hopf bifurcation in the Total Synaptic Coupling versus Noise Intensity for external bias current equal to 0.5 nA (full line) and 0.6 nA (dashed line) for hyperpolarizing (blue), shunting (red) and uniform distribution (green). For synaptic coupling higher than 250 nS (for 0.6 pA of bias current) and 300 nS (for 0.5 pA), a uniform distribution in synaptic reversal potential is less robust than shunting reversal potential.

## Discussion (1500 word limit)

### Type 2 Networks with Current Based Synapses

The transitions in Figs. 1 and 3 cross the border between asynchrony and synchrony to the stochastic population oscillator regime and to the coupled oscillator regime, respectively, and are equally well predicted by mean field theory (Fig. 4E). Another transition does not involve a bifurcation but instead involves a gradual increase in cycle skipping by which coupled oscillator neurons that fire early in a cycle suppress some of the other neurons on that cycle (Fig. 2), until the inhibition is so strong that neurons participate only sparsely and apparently randomly. A somewhat counterintuitive finding is that in “mean-driven” coupled oscillator regimes, the mean current can be subthreshold (Fig. 2A). The bifurcations and gradual transitions observed in networks of inhibitory neurons with type 2 excitability parallel those in networks with type 1 excitability (Brunel and Hansel 2006).

### Networks with Conductance Based Synapses

The transitions described in the preceding paragraph for current based synapses generalize to conductance based synapses. The representation of an inhibitory synapse by an outward current waveform implicitly as-sumes a hyperpolarizing synapse. Using conductance based synapses introduces an additional parameter, that of the synaptic reversal potential, which is neglected in the studies with current based synapses. Although shunting synapses do not carry the hyperpolarizing currents upon which the stochastic population oscillator theory is based, networks with shunting synapses nonetheless exhibited stochastic population oscillations (Fig. 6C). Moreover, the synaptic reversal potential parameter E_GABA,A_ exerts a powerful influence on synchronizing tendencies. In coupled oscillator networks, we have shown that the phase response curves of experimentally calibrated models of PV+ interneurons explain why hyperpolarizing synapses are more strongly synchronizing than shunting synapses (Via et al. 2022) (provided there are synaptic delays in the physiological range of 1 ms), in contrast to a previous study (Vida et al. 2006). The present study extends these results that hyperpolarizing synapses are more strongly synchronizing to the stochastic population oscillator regime.

The blue Hopf bifurcation curves in Fig. 7B for hyperpolarizing synapses have an inflection point at which the slope flattens out. We suspect that this is due to the saturating effect of hyperpolarizing conductance-based synapses. Whereas outward current synapses can cause an unlimited amount of hyperpolarization, conductance-based synapses cannot hyperpolarize the membrane potential beyond the reversal potential (-75 mV here), and therefore saturate. The green curves for a uniform distribution of firing rates also have an inflection point, but it occurs at higher values of conductance strength and is not as pronounced, likely due to the presence of shunting synapses as well as hyperpolarizing. The red curves for shunting inhibition (-55 mV) do not have an inflection point in the regime studied and have a slope intermediate between the steep and flat portions of the green curve, which results in the intersection points in which robustness of these two synaptic distributions to noise is reversed. At high levels of bias current, the membrane potential of individual neurons in the model may become sufficiently depolarized that synapses with a reversal potential of -55 mV have a small hyperpolarizing effect, which could be a potentially nonphysiological confound in our analysis.

### Heterogeneity

Here we have introduced temporal heterogeneity in that individual neurons receive distinct current noise. Actual inhibitory interneurons are heterogeneous in the synaptic connectivity, their passive and active properties (Fernandez et al. 2022; Via et al. 2022) and in their morphologies (Kriener et al. 2022). These types of heterogeneity are beyond the scope of this theoretical study.

### Increasing Gamma Synchrony as Therapeutic Strategy

PV+ fast spiking inhibitory interneurons have been shown to mediate gamma synchrony (Bartos et al. 2007; Cardin et al. 2009; Sohal 2022). In some cases, the gamma oscillations can be supported by inhibitory interneurons alone (Vinck et al. 2013, 2015, 2016) as in the computational examples in the current study, although there is frequently an interplay between inhibitory and excitatory cells (Tiesinga and Sejnowski 2009). There is mounting evidence to suggest that enhancing gamma synchrony may have multiple therapeutic benefits. Optogenetic stimulation of entorhinal cortex perforant path engram cells at high-gamma (100 Hz) frequency rescued memory impairments in a mouse model of AD (Roy et al. 2016). Moreover, gamma oscillations can also affect molecular pathology; entrainment of gamma in the visual cortex by 40 Hz flickering light and by optogenetic stimulation of PV+ interneurons in hippocampal area CA1 in a mouse model of AD reduces Aβ levels in those respective areas (Iaccarino et al. 2016).

In a coupled oscillator regime, enhancing gamma synchrony could involve targeting specific intrinsic ion channels to fine tune the phase resetting properties. Regardless of the oscillatory mechanism, this study shows that the reversal potential for GABA^A^ receptor chloride channels is a likely therapeutic target to enhance gamma synchrony. However, it seems likely that the synaptic reversal potential is heterogeneous across synapses (Blaesse et al. 2009; Otsu et al. 2020). If we assume a physiological network with the synaptic reversal potential distributed uniformly between the two extreme values, then manipulations to reduce internal Cl^-^ concentrations in PV+ interneurons should increase gamma synchrony, because networks with purely hyperpolarizing synapses synchronize more robustly than those with a uniform distribution (Fig. 7).

### Implementation of Therapeutic Strategies

In mature neurons, the reversal potential of the GABA_A_ synapses is thought to be determined by the expression of anion cotransporters, primarily KCC2 (Blaesse et al. 2009). However, bicarbonate ions also flow through GABA_A_ channels, causing the reversal potential of GABA_A_ to be more depolarized than the reversal potential for chloride E_Cl_ (Blaesse et al. 2009). One caveat in designing therapies to modify E_Cl_ is that chloride microdomains due to the inhomogeneous distribution of anionic polymers such as actin, tubulin, and nucleic acids (Rahmati et al. 2021) may exert a strong local influence on E_Cl_. There is growing evidence that a defective ratio of chloride importer NKCC1 and chloride exporter KCC2 is present in several neurodevelopmental conditions (Savardi et al. 2021). Since these cotransporters are fundamental in the regulation of neuronal chloride concentration, therapies focused on restoring chloride homeostasis, and hence modifying E_GABA,A_, could have an impact in restoring core symptoms of several disorders. For example, (Parrini et al. 2021) showed that reducing NKCC1 expression in a Ts65Dn mouse model of Down syndrome restores the intracellular chloride concentration, efficacy of GABA-mediated inhibition, and neuronal networks dynamics, rescuing cognitive deficits as well. Thus, although there are many technical challenges to selectively targeting the reversal potential for GABA_A_ synapses selectively in PV+ inhibitory interneurons, this strategy could have powerful implications for gamma synchrony and its potential role in reversing cognitive impairment.

## Acknowledgements

This work was funded by NIH NS054281 to CCC and utilized resources provided by NSF 2018936. We thank Nicolas Brunel for comments on an earlier draft of this manuscript.

